# The organization and developmental establishment of cortical interneuron presynaptic circuits

**DOI:** 10.1101/2020.09.17.302117

**Authors:** Gabrielle Pouchelon, Yannick Bollmann, Elaine Fisher, Chimuanya K Agba, Qing Xu, Kimberly D Ritola, Andrea MC Mirow, Sehyun Kim, Rosa Cossart, Gord Fishell

**Affiliations:** Harvard Medical School, Department of Neurobiology, Boston, Massachusetts 02115, USA; Broad Institute, Stanley Center for Psychiatric Research, Cambridge, Massachusetts 02142, USA; Aix Marseille University, INSERM, INMED, Turing Center for Living Systems, Marseille, France; Janelia Research Campus, Howard Hughes Medical Institute, Ashburn, VA 20147, USA; Center for Genomics & Systems Biology, New York University, Abu Dhabi, UAE

## Abstract

Sensory and cognitive functions are processed in discrete cortical areas and depend upon the integration of long range cortical and subcortical inputs. PV and SST inhibitory interneurons (cINs) gate these inputs and failure to do so properly is implicated in many neurodevelopmental disorders. The logic by which these interneuron populations are integrated into cortical circuits and how these vary across sensory versus associative cortical areas is unknown. To answer this question, we began by surveying the breadth of afferents impinging upon PV and SST cINs within distinct cortical areas. We found that presynaptic inputs to both cIN populations are similar and primarily dictated by their areal location. By contrast, the timing of when they receive these afferents is cell-type specific. In sensory regions, both SST and PV cINs initially receive thalamocortical first order inputs. While by adulthood PV cINs remain heavily skewed towards first order inputs, SST cINs receive an equal balance of first and higher order thalamic afferents. Remarkably, while perturbations to sensory experience affect PV cIN thalamocortical connectivity, SST cIN connectivity is disrupted in a model of fragile X syndrome (Fmr1 loss of function) but not a model of ASD (Shank3B loss of function). Altogether, these data provide a comprehensive map of cIN afferents within different functional cortical areas and reveal the region-specific logic by which PV and SST cIN circuits are established.

---

Our conscious perception of the world is rooted in the neocortex. Integration of sensory information and the subsequent generation of higher cognitive functions, such as arousal, attention or prediction^1,2^, are processed in discrete cortical areas. Across the cortex, GABAergic PV and SST cells, the two largest classes of cortical interneurons (cINs), form local computational units that gate sensory, modulatory and intercortical information ^3^. Recent work has shown that at a genetic level each of the discrete cIN subtypes is remarkably similar regardless of cortical area^4^. Combined with circuit analysis^5,6^, this has led to the suggestion that PV and SST cINs utilize common motifs irrespective of the cortical circuits in which they are imbedded. This is puzzling given the functional differences across sensory, motor and associative areas. Perhaps, while the local connectivity of PV and SST cINs is shared across regions, their functional engagement is specialized through region-specific differences in their afferents. To clarify this issue, we undertook a systematic examination of the afferent connectivity of PV and SST cINs in two sensory (S1 and V1) and one associative (M2) regions. To do so, we did a broad-scale mapping of presynaptic inputs using monosynaptic rabies tracing^7,8^. This revealed that while within specific regions PV and SST cINs receive similar afferents, these vary in accordance with the cortical area examined.

It is well accepted that inputs play a critical role in the development of cortical neurons^9-14^. Although PV and SST cINs in specific regions share common inputs, we wondered whether the developmental timing when they receive these afferents differ. To explore this question, we examined when specific afferent populations innervate these two cIN classes during development. This revealed that depending on region, afferents onto PV and SST cINs differentially acquire permanent and transient afferents in distinct temporal orders.

cIN dysfunctions have been described in many neurodevelopmental disorders to result from both environmental and genetic insults. Understanding of how such abnormalities arise has come from work demonstrating that the differentiation of cINs is controlled by both sensory activity^15,16^ and intrinsic genetic programs ^17-19^. Although varying in accordance with the cortical region examined, the thalamus provides the principal sensory relay onto PV and SST cINs. The thalamus has two major types of cortical efferents. From the First-Order neurons (FO) ^20^, cINs receive “passive” information ^21,22^, while the Higher-Order (HO) neurons, by integrating multimodal information, provide cINs “active” sensory input ^23-26^. We found that PV and SST cINs in V1 and S1 receive different proportions of FO versus HO inputs. While SST cINs receive equivalent amounts of FO/HO afferents, PV cINs are primarily driven by FO thalamic input. To examine the importance of sensory input to the development of PV versus SST cINs, we disrupted somatosensory and visual activity during development. Within PV cINs this resulted in these cells receiving supernumerary HO inputs. Surprisingly FO/HO balance onto SST cINs was not affected. We therefore hypothesized within SST cINs the ratio of FO/HO input is genetically controlled. We examined this question in two monogenic models of disease, *Shank3b*^27^ and *Fmr1* KOs ^28,29^.While the ratio of FO/HO inputs was unaffected in *Shank3b* mutants, in *Fmr1* nulls, this ratio was strongly shifted in favor of FO inputs, such that it resembled that seen in early development.

Altogether, these data provide a comprehensive map of cIN afferents within different functional cortical areas. This revealed the region-specific logic by which PV and SST cIN circuits are established. Finally, we discovered that while PV cIN connectivity in sensory areas is mediated by activity, SST cIN circuits are predominantly genetically determined.

## Results

### Mapping developmental changes in the afferent connectivity of PV and SST interneurons

In order to understand the areal differences in connectivity of PV and SST cINs within infragranular cortex across development, we restricted our analysis to layers 5 and 6 of either primary sensory (S1 and V1) or associate (M2) areas (Extended Data Fig. 1 a). To determine the breadth of presynaptic connectivity impinging upon these populations, we utilized a genetically modified form of CVS N2c (N2cRV) rabies^8^. This strain is less toxic and more comprehensively reports afferent connectivity than the B19 variant originally utilized ^7^. Rabies tracing is achieved by infecting starter cells with an AAV-helper that provides dual complementation allowing for rabies infection (TVA) and monosynaptic transport (G protein) (Fig. 1a). Both helper elements and a reporter (eGFP) were combined into a single AAV-helper virus (AAV-DIO-TGN, Fig. 1b). This was essential for accurate determination of starter cells and quantification of connectivity.

**Fig. 1:**
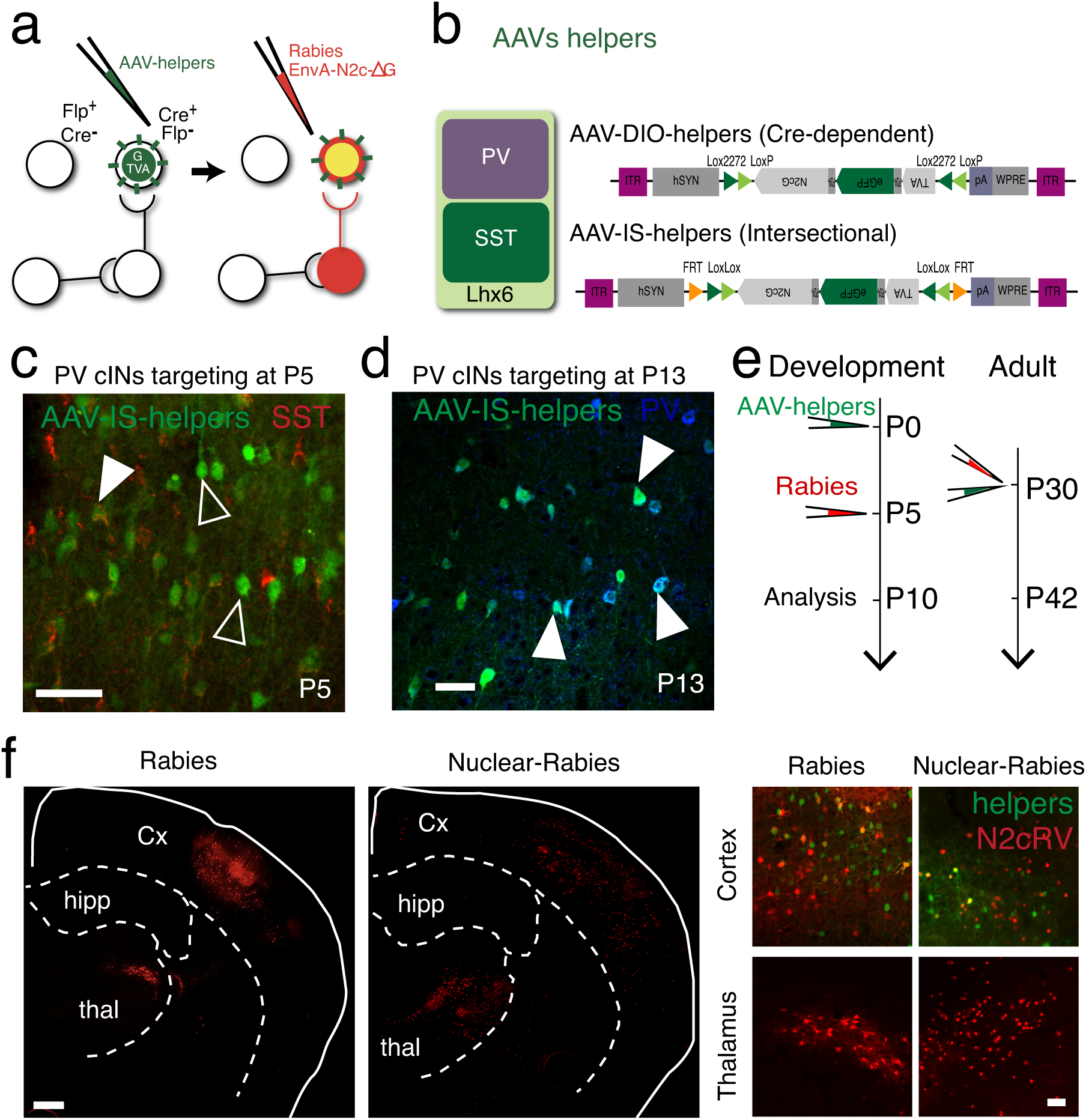
Mapping developmental changes in the afferent connectivity of PV and SST cINs. **a**, Schematic of the principle for monosynaptic rabies tracing. TVA and G conditional helpers (in green) are expressed using AAVs, followed by the specific infection and retrograde labeling by EnVA-pseudotyped CVS N2c rabies virus (N2cRV, in red). **b**, PVcre- and SST-cre mouse lines are used with AAV-DIO-helpers construct (top). Intersectional strategy using Lhx6-iCre/SST-FlpO mouse lines is used with the AAV-IS-helper (bottom). **c**, Somatostatin staining (red) is absent from AAV-IS helpers infected cells (green) within Lhx6iCre/SSTFlpO mice showing the PV cIN specificity as early as P5. (scale bar 50μm). **d**, Parvalbumin staining (blue) showing the PV cIN identity of AAV-IS helpers infected cells (green) at P13 (scale bar 50μm). **e**, Timeline of AAV-helpers and N2cRV injections for the developmental timepoint (left) and for the adult timepoints (right). **f**, Example of retrograde-labeling using Rabies virus expressing etierh an mCherry reporter (left) or a nuclear-tdtomato (right) (scale bar 500μm). The insets from the cortex (top) and the thalamus (bottom) show the better cellular detection with the nuclear expression (scale bar 100μm). **g**, 3D representation of N2cRV tracing examples from the 3 distinct areas targeted (M2, S1, V1) and from PV or SST cINs) using Brainrender^86^.

We used SST-Cre mice in order to target SST cINs for both adult and development. This was possible as somatostatin expression initiates sufficiently early to be used perinatally. However, while we could use PV-Cre to target adult PV cINs, the late onset of parvalbumin gene expression prevents its use for developmental time points (up to P20 in V1. Extended Data Fig. 1c). For their early targeting, we therefore developed a Boolean-based intersectional AAV strategy ^30^. Lhx6^+^ progenitors give rise to both SST and PV cINs and provide an early marker for both populations. The coincident early expression of somatostatin with Lhx6 allowed us to implement a subtractive (Lhx6-iCre-ON/SST-FlpO-OFF) strategy for selectively targeting early PV cIN populations. Lhx6-iCre triggers the expression of DIO helpers, while SST-FlpO abrogates the expression of this virus within the SST population (AAV-IS-helpers, Fig. 1b). While the AAV-helper virus was injected at P0, N2cRV was injected at P5. This was necessary to allow for the suppression of helper virus within the off-target SST population (Fig. 1e). To verify the PV targeting specificity, we confirmed that the targeted population was uniformly SST-negative. Five days post-injection, we found a high specificity for putative PV cINs (Fig. 1c; Extended Data Fig. 1d; 73.67% ±3.03 in M2; 73.77%±0.84 in S1; 76.60±3.96 in V1). We also confirmed that this population later expressed parvalbumin (Fig. 1d).

The degree of connectivity of specific PV or SST starter cells was normalized in accordance with the relative percentage of afferents from different structures. Secondarily, we also compared the relative numbers of afferents from a given structure that target either PV or SST cINs (Extended Data Fig. 1b, n=21 adult control animals (n=11/10 PV/SST cINs; n=3-4 per area and n= 20 control P10 animals; n=9/11 PV/SST cINs; n=3-4 per area). In addition, we created a N2c rabies expressing a nuclear tdtomato reporter (NLS:tdtomato, Fig. 1f), which improved our ability to accurately access afferent cell number.

### Presynaptic inputs to PV and SST cINs differ in accordance with cortical location

Previous studies only detected very small differences between the connectivity of cIN subtypes using rabies tracing but did not compare their connectivity within different cortical areas ^31-34^. We hypothesized that differences in connectivity rely upon the regional location in which PV and SST cells settle, as suggested previously from medial prefrontal cortex (mPFC) tracing^35^. We therefore examined PV and SST cIN afferent connectivity across three distinct cortical areas: two sensory areas, primary visual (V1) and somatosensory (S1) and a non-sensory associative area, the premotor cortex (M2) (Fig. 1e). We found that the number of local projection neurons of both PV and SST cINs within M2 was lower than that observed within V1 and S1 areas (Fig. 2a; Extended Data 3a; one-way ANOVA p<0.01). In contrast, long range cortical connectivity (contralateral neurons and other cortical areas) to M2 cINs was higher (Extended Data Fig.2a, Fig3b;). Interestingly, the identity of neurons projecting from other cortical areas is specific to their target (Fig. 2b; one-way ANOVA p<0.0.01). For instance, retrosplenial cortex projects to V1 cINs, while cingulate projects to S1 and M2 cINs (Fig. 2b). M2 cINs also receive more subcortical modulatory inputs (Extended Data Fig. 2c-d), such as cholinergic basal forebrain inputs (Fig. 2d; one-way ANOVA p<0.0001; Extended Data Fig. 2e), as well as inputs from serotoninergic mid/hindbrain regions (Fig. 2e; one-way ANOVA p<0.001; Extended Data Fig. 2e). In addition, M2 cINs receive input from the claustrum, while in sensory regions they do not (Extended Data Fig. 2b). Pearson’s correlation calculated using all afferent structures (correlation matrix Fig. 2f, degree of connectivity heatmap Fig. 2f), as well as the network created using Fruchterman-Reingold algorithm (Extended Data Fig. 2f) show that the afferent connectivity of PV and SST cINs from different cortical regions is distinct, while PV and SST cINs from a given cortical area group together (Fig. 2f) Therefore, adulthood the presynaptic inputs to SST and PV cINs in adulthood are reflective of their regional position not their subtype identity.

**Fig. 2:**
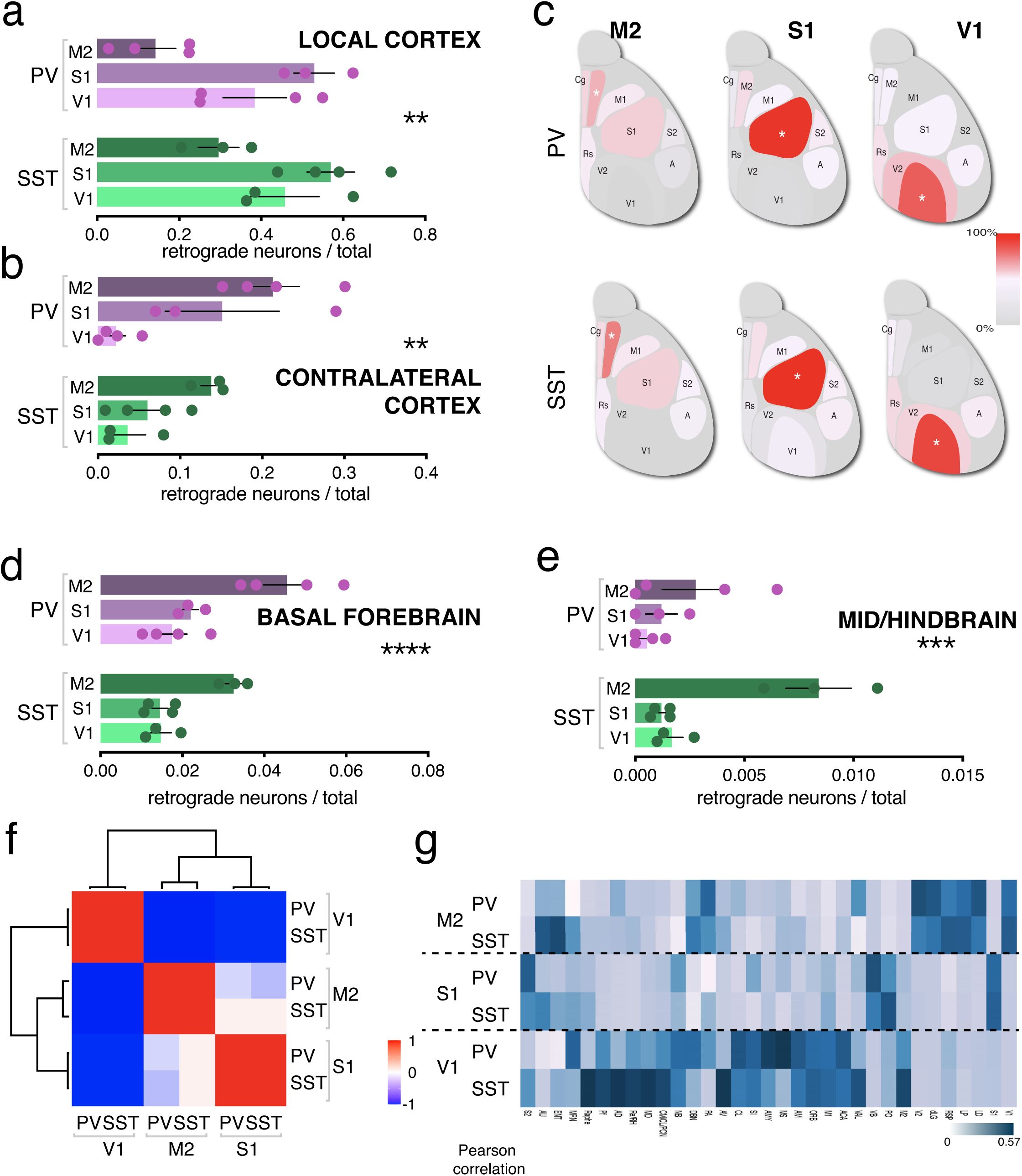
Presynaptic inputs to PV and SST cINs are specific to their cortical areal location. **a**,**b**, The degree of local (**a**) and contralateral (**b**) retrogradely labeled neurons to PV (purple) and SST (green) cINs within M2, S1 and V1 shows areal differences of connectivity (one-way ANOVA p<0.01), where both PV and SST cINs in M2 show lower local, but higher contralateral connectivity compared to sensory areas. **c**, Heatmap showing the percentage of connectivity within ipsilateral cortical afferent neurons to PV and SST cINs traced from M2, S1 or V1. The heatmap show (n=21, averaged per condition). **d, e**, The degree of connectivity of basal forebrain(d) and midbrain and hindbrain (e) were both higher to PV and SST cINs in M2 than cINs in sensory areas (one-way ANOVA p<0.0.01). **f**, Heatmap representing Pearson’s correlation coefficient between PV and SST cINs in M2, S1 and V1 using the average of the degree of connectivity for all structures labeled in each case, showing that PV and SST cINs are highly correlated when located in a same area, M2, S1 or V1. **g**, Heatmap representing the degree connectivity average (n=21) for afferent structures to PV and SST cINs in M2, S1 and V1.

### During development presynaptic inputs to PV and SST cINs are cell-type specific and dynamically regulated

Although in PV and SST cINs presynaptic inputs are anatomically similar at mature stages, they are functionally extremely different. PV cINs are directly involved in controlling sensory inputs through feedforward inhibition^36^, while SST cINs utilize feedback inhibition^37^. They are also differentially involved in cortical processing, such as fear memory in mPFC ^38^, oscillations in V1^39^ and whisker-dependent touch in S1^26^. It is well accepted that timing is essential for their function in adult learning^40^, as well as their proper integration into cortical circuits ^41^. Therefore, we wanted to examine how this principle influences cIN postnatal development. Specifically, we hypothesized that the mature functional differences in PV and SST cINs arise from their dynamics in the establishment of afferent connectivity. Indeed, when we compared the mature connectivity of these populations to that during development, we found cell-type specific differences. We observed two trends: a precise temporal ordering as to when specific projections are established and transient connectivity where afferent regions that are initially highly connected are strongly reduced or lost at more mature stages (Extended Data Fig. 3,4). Notably, these changes occur in a region-specific manner. For example, although in mature M2 cortex both PV and SST cINs have low local connectivity (see above, Fig. 2a), PV cINs at early postnatal times are highly connected while SST cINs are not (Fig. 3a; Student t-test p<0.05 PV cINs in dev vs adult). Consistent with the differential dynamics in the development of PV versus SST cINs within M2, while both populations receive substantial long-range cortical connectivity in adults, perinatally PV cINs have few such inputs. By contrast, SST cINs already have strong long-range connectivity at early postnatal times (contralateral Fig. 3a; Student t-test p<0.01; ipsilateral, Extended Data Fig. 3c).

**Fig. 3:**
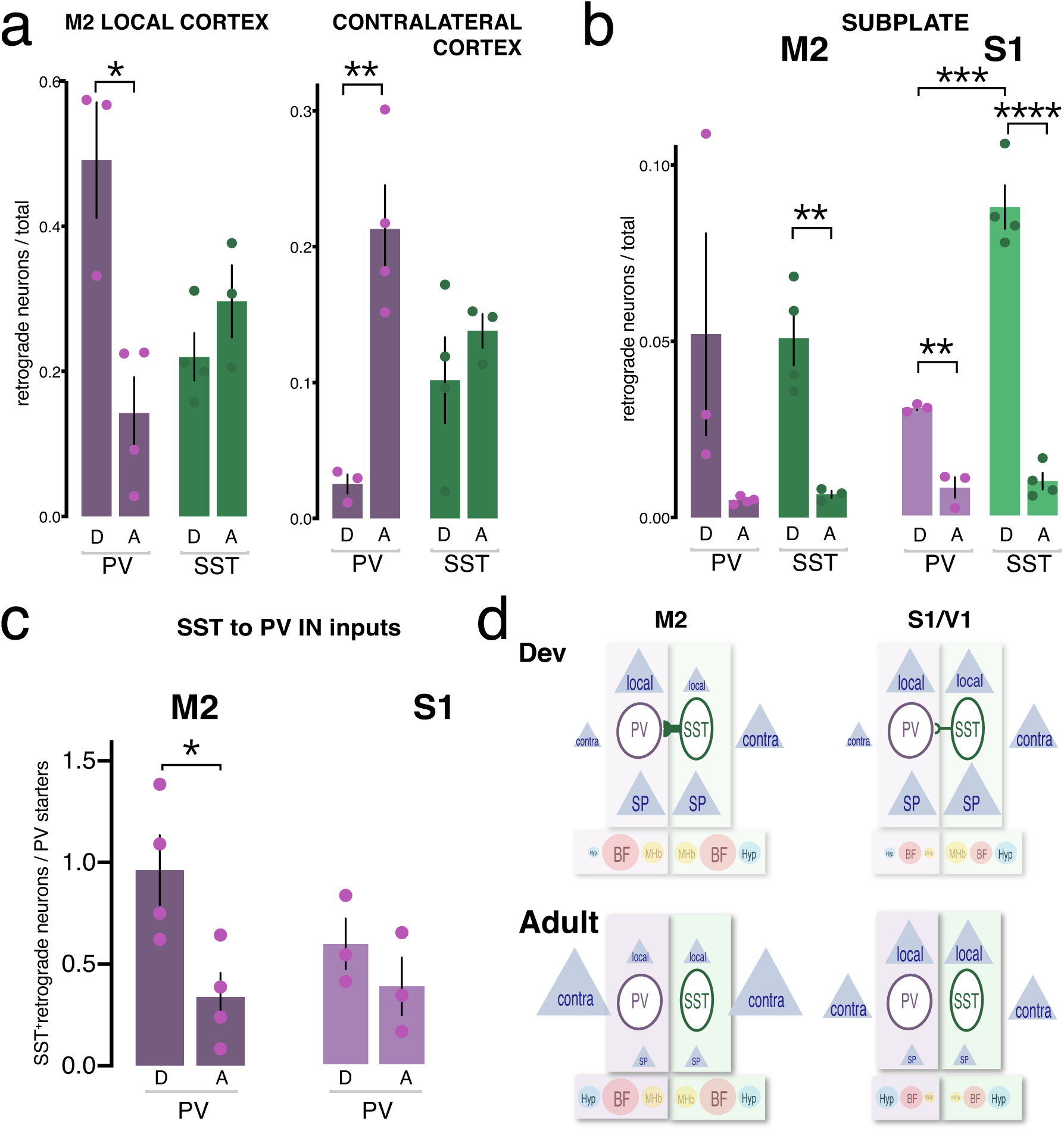
During development, presynaptic inputs to PV and SST cINs are cell-type specific and dynamically regulated. **a**, Local connectivity (left) is higher, while contralateral connectivity (right) is lower onto PV cINs (purple) in M2 during development (Student t-test p<0.01 PV cINs in dev vs adult). In contrast, both connectivity types onto SST cINs are consistent between adult and development. **b**, Subplate transient neuron projections are higher to both PV and SST cINs during development (Student t-test SST cINs dev vs adult in M2 PV p<0.01; PV cINs dev vs adult in S1 SST p<0.01; SST cINs dev vs adult in S1 p<0.0001). However, subplate connectivity is higher onto SST than PV cINs in S1 (Student t-test p<0.001). **c**, SST cIN transient connectivity to PV cINs is higher inM2 (Student t-test p<0.05), but not in S1. **d**, Diagram summarizing all the types of afferents received by SST and PV cINs within M2 and sensory areas (S1/V1) during development and at adulthood. Left side of each diagram represents PV cIN connectivity, while the right side represents SST cIN connectivity. *D or Dev = development; A= adult; Hyp = hypothalamus; BF = basal forebrain; MHb=Midbrain and Hindbrain; SP = supblate; contra = contralateral*.

**Fig. 4:**
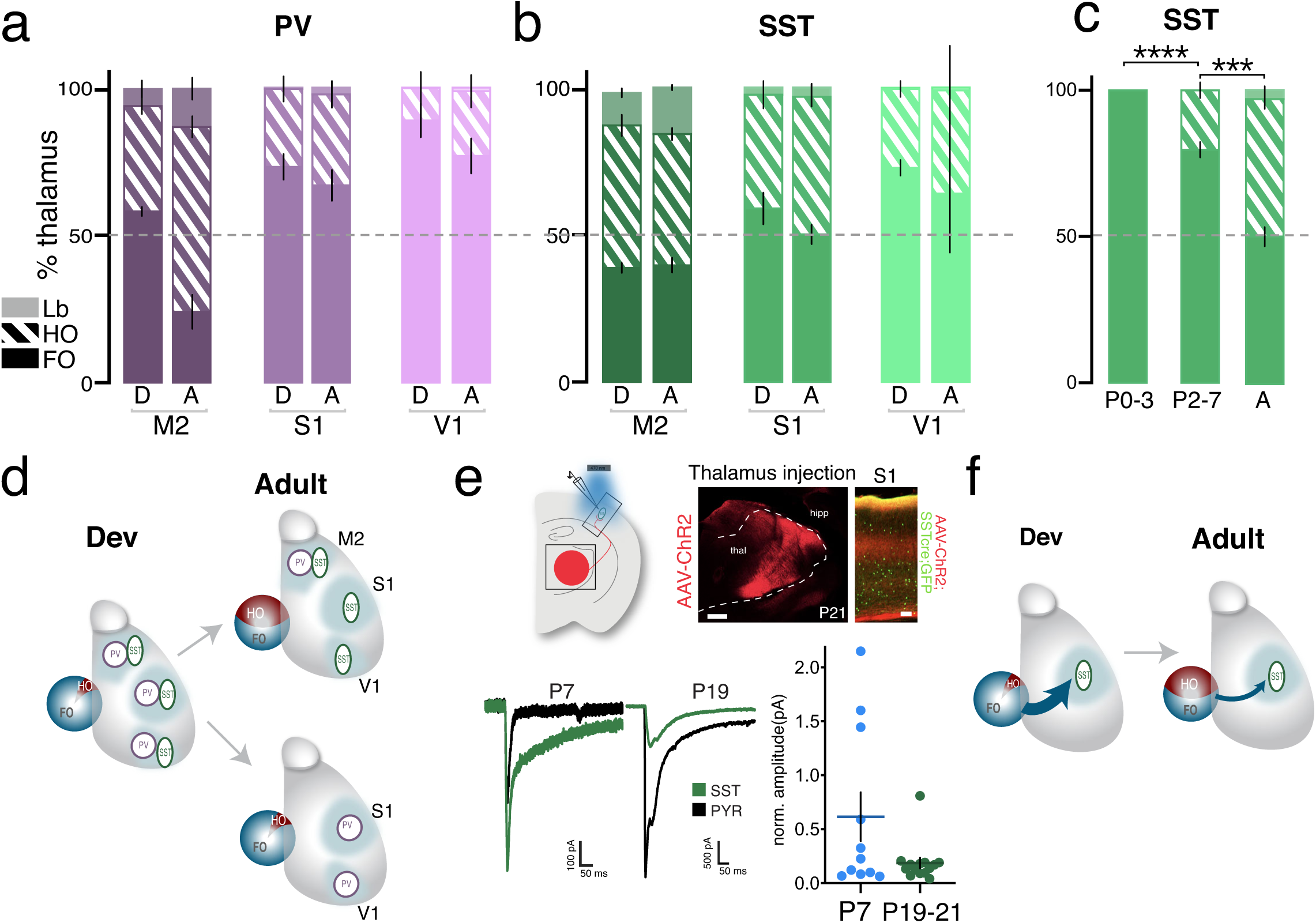
Thalamocortical afferents to SST and PV cINs are developmentally regulated in a regional and cell-type specific manner. **a, b**, Ratio of FO, HO and Limbic neurons within the whole thalamus (100%) retrogradely labeled with N2cRV. In contrast to M2, PV cINs in S1/V1 receive higher proportion of FO afferents both during development (P5-P10) and in adult (**a**). SST cINs receive a higher proportion of HO afferents to than PV cINs in sensory areas (S1 and V1). SST cINs tend to have more FO during development (**b**). **c**, Early developmental dynamics of thalamocortical afferents onto SST cINs in S1. Compared to adult control (from b), SST cINs receive more FO afferents than HO at P2-P7 (Student t-test p<0.001) and only FO afferents at P0-P3 (from b; Student t-test p<0.0001). **d**, Diagram summarizing the change in the ratio of FO and HO thalamic afferents onto PV and SST cINs in development and in adult. **e**, Optogenetic response of SST cINs to thalamic afferent stimulation (left). AAV-ChR2 was injected in the thalamus (AAV-ChR2 in red, scale bar 500μm) and SST cINs labeled with a Cre-dependent reporter a cortex (green, scale bar 100μm), together with negative pyramidal cells (black) were recorded in response to light stimulation. At P7, SST cIN response amplitude normalized by pyramidal cell amplitude, is higher than in mature stage P19-P20. **f**, Summary diagram including both the dynamic of FO/HO ratio and the thalamocortical strength over development of SST cINs in S1.

Similarly, transient connectivity to PV versus SST cINs was distinct in specific cortical regions. To investigate this, we quantified subplate neurons, and confirmed their subplate identity using CTGF^+^ staining^42^. While subplate connectivity is transient on both PV and SST cINS (S1 dev vs adult PV p<0.01; SST p<0.0001), S1 SST cINs have considerably higher afferent connectivity than either PV cINs or SST cINs within M2 at P10. (Fig. 4cb; p<0.001). We and others have previously shown that in S1, PV cINs receive strong transient inputs from SST cINs ^12,43^ that control their maturation. Following N2cRV tracing of PV cINs, we stained for somatostatin and found that connectivity was higher during development onto M2 PV cINs (Fig. 3c, Student t-test PV cINs in M2 dev vs adult p<0.05), while it was not in S1. Taken together, while the ultimate connectivity of both populations is similar in the adult, the developmental dynamics by which this is achieved are markedly different both within these two populations and across regions (Fig. 3d; Extended Data Fig. 3,4).

### Thalamocortical afferents to SST and PV cINs are regional and cell-type specific and are developmentally regulated

Sensory experience transmitted through the thalamus is critical for the development of cortical areas ^44-47^. The thalamus is organized into anatomically distinct nuclei that possess specific functions, broadly classified as limbic, First-Order (FO) and Higher-Order (HO). We examined the identity of TC neurons projecting to PV and SST cINs from each of these divisions in sensory and associative cortical areas. Recent studies have enabled the genetic identification and classification of thalamic nuclei^48,49^ (Extended Data Fig. 5a). This allowed us to anatomically group thalamic neurons retrogradely labeled from PV and SST cINs into these 3 functional categories. For example, V1 and S1 cortices receive both FO and HO afferents, from dLG/LP and VB/PO, respectively (Extended Data Fig. 5b). We examined the percentage of FO, HO and limbic TC neurons projecting to PV and SST cINs in each cortical area and found that PV cINs receive more FO projections than SST cINs in both V1 (mean±SEM PV: 76.88±5.90 vs SST: 64.03±17.64%) and S1 (mean±SEM PV: 66.93±5.220 vs SST: 50.13±3.256%), while in both sensory areas the proportion of limbic inputs was negligible. By contrast and as expected, in M2 both PV and SST cINs receive primarily HO (mean±SEM PV: 62.68±3.60; SST: 44.52±2.05%), as well as limbic (mean±SEM PV: 12.92±3.70; SST: 15.79±0.845) afferents (Extended Data Fig. 5d). We next looked at the developmental dynamics by which TC hierarchy is established. We found that both subtypes first receive FO inputs from early development onwards (Extended Data Fig. 5c,d) but only later obtain HO afferents. Notably, PV cINs in S1 and V1 continue to acquire FO afferents throughout their maturation. Although at P0, SST cINs in sensory areas only receive FO afferents (Fig. 4c P3 vs P7 p<0.0001), as development progresses, they acquire ever greater proportions of HO afferents (P7 vs adult p<0.001). This results in them having equal numbers of FO and HO afferents by adulthood (Fig. 5b &c; Extended Data Fig. 5c, d). Perhaps because the rabies labeling window for adult brains overlapped with the visual critical period (P30-P42), the variability was higher in V1.

**Fig. 5:**
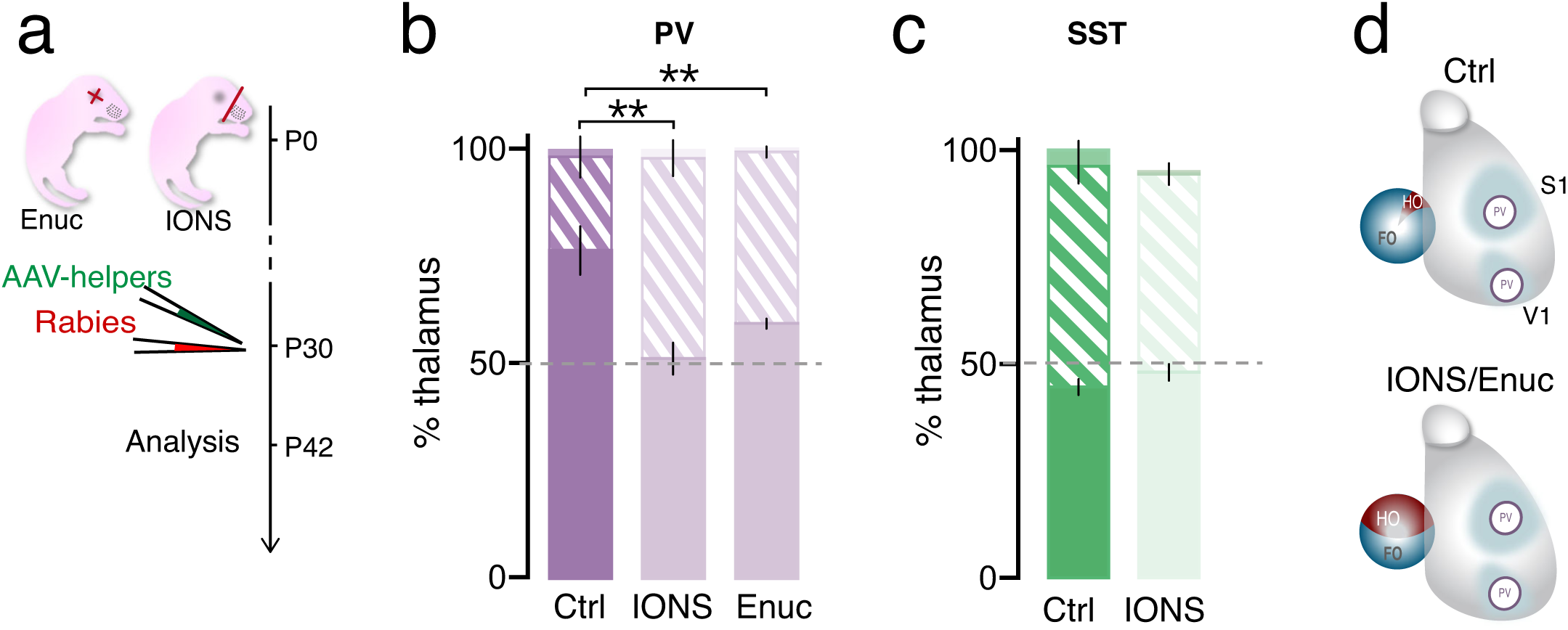
The ratio of FO/HO thalamocortical projections onto PV cINs is experience dependent. **a**, Experimental timeline: with sensory deprivation of the visual system (Enuc = enucleation) or of the somatosensory system (IONS = infraorbital nerve section) at P0, followed at P30 by the rabies virus (N2cRV) and AAV-helpers injection. **b**, PV cINs receive more FO projections than SST cINs in both V1 and S1 (mean±SEM Ctrl 76.60±5.63% vs IONS 51.38±3.72% vs Enuc 59.53±1.15%; n= 3/4/3; Student t-test p<0.01). **c**, The ratio of FO/HO projections onto SST cINs in S1 is unchanged upon IONS (mean±SEM Ctrl 44.64±1.81% vs IONS 48.07±1.92%; n=3/3). **d**, Summary diagram of Ctrl and deprived thalamocortical afferent ratio.

FO and HO neurons transmit different types of information. We therefore investigated whether the difference in input proportions onto SST cINs also reflects differences in TC input strength. To do so, we stereotaxically injected an AAV-expressing hChr2 in the thalamus and recorded SST cINs at both mature (P21) and postnatally (P7) time points. As previously shown^43^, in comparison to pyramidal neurons the relative amplitude of responses decreases during maturation (Fig. 4e; mean±SEM P7: 0.6158±0.43, P21: 0.1867±0.21). This confirms that the decrease in strength of thalamic inputs onto SST cINs across development occurs concomitant with increases in their receipt of HO TC afferents (Fig. 4d & f).

### The ratio of FO/HO thalamocortical projections onto PV cINs is experience dependent

The hierarchy of TC pathways onto PV and SST cINs is dynamically regulated during the postnatal stages when early sensory activity is essential to shape circuitry in visual^50,51^ and somatosensory areas^43,44^. We therefore tested whether early sensory activity controls the hierarchy of TC pathways onto PV and SST cINs. We performed either infraorbital nerve section (IONS) or enucleation (Enuc) at P0 to disrupt whisker-dependent and visual inputs, respectively. We then used rabies to retrogradely label thalamic afferents to PV and SST cINs from S1 or V1 and quantified the ratio of mature FO and HO neuron projections. We found that the normal hierarchy onto PV cINs but not onto SST cINs was disrupted (Fig. 5a; Extended Data Fig. 6). In both enucleated and whisker-deprived animals, the FO/HO ratio was decreased so that it resembled that seen onto SST cINs (mean±SEM Ctrl 76.60±5.63% vs IONS 51.38±3.72 vs Enuc 59.53±1.15; n= 10; Student t-test p<0.01). By contrast, the FO/HO ratio within SST cINs was unchanged in sensory deprived animals. This suggests that during development, the TC afferents to PV cINs is more plastic than to SST populations. This may reflect the strong FO sensory inputs that are maintained within PV cINs across development.

**Fig. 6:**
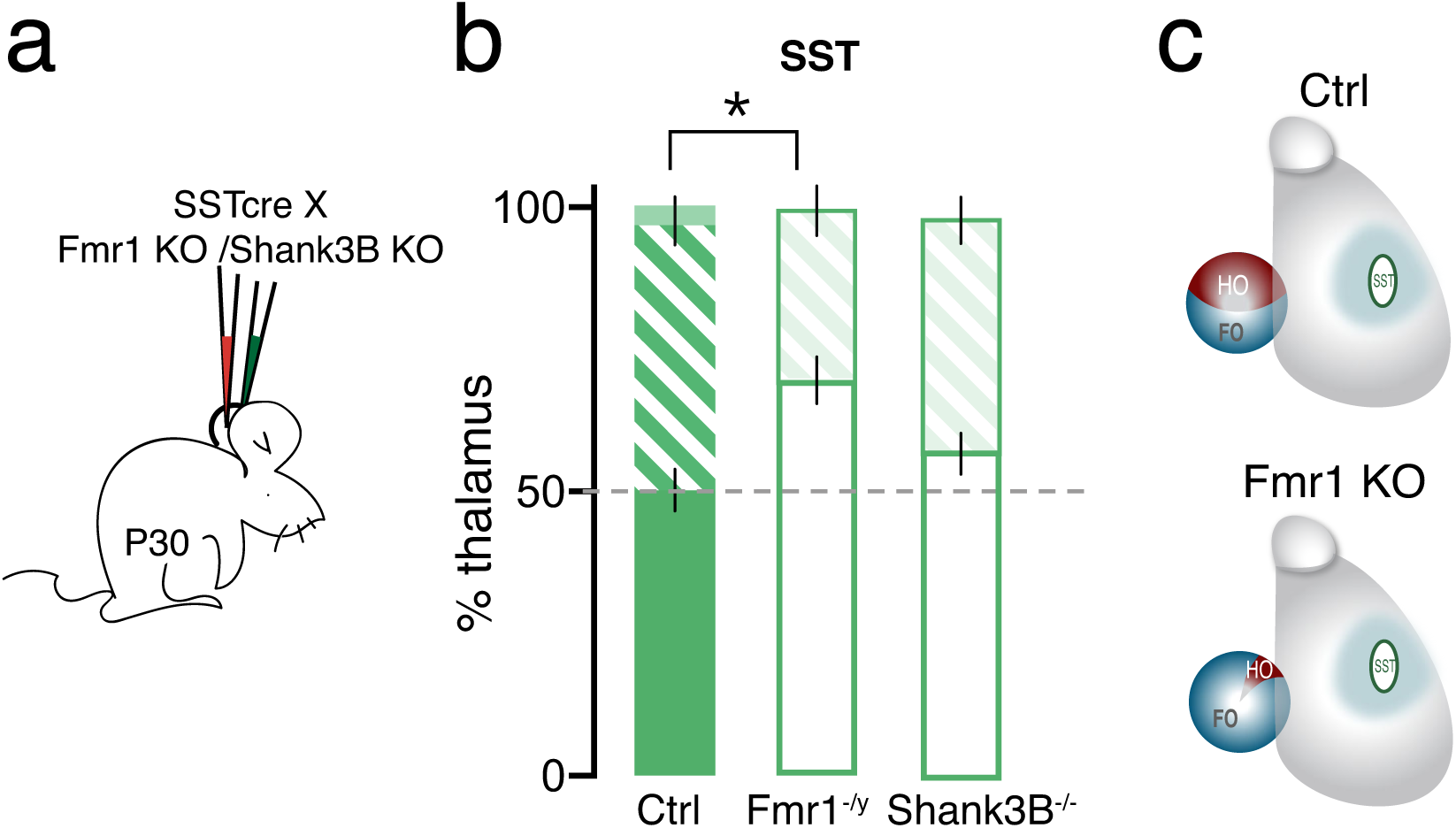
The ratio of FO/HO thalamocortical projections onto SST cINs is genetically regulated. **a**, FO/HO ratio within the whole thalamus (100%) retrogradely labeled with N2cRV in neurodevelopmental disorder models: autism spectrum disorder with *Shank3B*-/- and Fragile X syndrome disorder with *Fmr1*-/y. **b**, FO/HO ratio is increased at the advantage of FO afferents in Fmr1-/y but not Shank3B-/- (mean±SEM Ctrl 51.98±7.42% ; n=5 (3 from Fig. 4b) vs Fmr1-/y 69.41±4.10%; n=5; Student t-test p<0.05; vs Shank3B 56.80±3.60%; n=4). **c**, Summary diagram of Ctrl and fmr1 -/y thalamocortical afferent ratio.

### The ratio of FO/HO thalamocortical projections onto SST cINs is genetically regulated

Dysfunction in both cINs and thalamocortical circuits has been implicated in neurodevelopmental conditions, such as schizophrenia and autism spectrum disorders (ASD) ^52,53^. Hypersensitivity and somatosensory behavioral defects in many neurodevelopmental disorders^54,55^ suggest they may involve both areal-specific and cell-type specific circuit defects. While in most cases these are known to have a strong genetic component, the precise causal mutations are both various and complex. We therefore explored two monogenetic neuropsychiatric models, *Fmr1* and *Shank3* LOF. These models are attractive as both have been shown to reproduce in mice the core phenotypes seen in humans lacking these genes ^29,56^. As shown above, disruptions in sensory signaling affect the ratio of FO to HO TC afferents in PV but not SST cINs. The absence of environmental impact on the ratio of FO to HO afferents onto SST cINs led us to hypothesize that this connectivity may genetically regulated. We therefore performed rabies tracing in S1 SST cINs in *Shank3B* null or male *Frm1* KO mice. We found that in *Fmr1* but not *Shank3B* null mice, the ratio of FO to HO TC afferents onto SST cINs was altered (Fig. 6b; FO% Ctrl ± 50.13±3.256 *vs* KO 69.60±4.11). Specifically, in *Fmr1* mice the number of FO neurons projecting to SST cINs was increased at the expense of HO afferents (Extended Data Fig. 7). Thus, in *Fmr1* mice the adult TC SST cIN connectivity resembled that seen in young animals (Fig. 6c).

## Discussion

In this paper we addressed the paradox that despite considerable function differences between regions, specific types of cINs form similar anatomical circuits^31-33,35^. Two central principles emerged: the regional location of cINs defines their presynaptic inputs, while the timing when their afferent input is established in specific regions is best predicted by cell type. Furthermore, while PV and SST cINs in all regions receive thalamic input, the types of input depend both on cell type and the region of cortex examined. PV cINs in sensory areas are unique in that they primarily receive FO afferents. However, while SST cINs also first receive mainly FO inputs, by adulthood they have equal numbers of FO and HO afferents. Within PV cINs the afferentation pattern was determined by sensory experience, while the organization of SST cINs was affected by genetic perturbations.

Understanding the developmental dynamics by which cIN connectivity is established is fundamental but difficult to explore. Optogenetic tools in the last decade have proved essential for the study of cell-type specific circuits^25,57-60^. Optogenetics however primarily allows for the detection of strong inputs, while rabies tracing reveals networks regardless of strength. As such, at present despite some concerns about specificity^61^ rabies remains the best tool available for surveying the connectome^62^. Importantly, in both our experience^13,43,63^ and others^64,35,65^, to date all monosynaptic connectivity detected by rabies can be confirmed by other methods. Albeit, a degree of tropism in rabies tracing does exist, as certain synaptically connected types appear resistant to rabies labeling^61^. One also cannot compare the strength of connectivity to the number of afferent neurons. Despite these caveats, within distinct regions of the cortex our rabies tracing accurately reveals both developmental increases and decreases in connectivity of PV and SST cINs.

Our study discovered the developmental appearance of neuromodulatory afferents to PV and SST cINs. For example, from early postnatal development we observed substantial cholinergic afferents, which are difficult to detect using optogenetic or electrical methods ^66,67^. In addition, we revealed the developmental timing when afferents from preoptic hypothalamus{K:wr}, VTA, subthalamic ^68 69^ and raphe inputs are established. Lastly, our data revealed these projections arise from a variety of neuronal subtypes. For instance, within the basal forebrain and raphe, in addition to cholinergic and serotoninergic subtypes, were unstained populations that are likely glutamatergic or GABAergic.

One of the more puzzling aspects of our findings is that despite receiving similar modulatory and cortical afferents, behavioral studies indicate that PV and SST cINs integrate them differentially^70-72^. This in part is likely achieved by the transcriptional differences between these cell types. This suggests that the postsynaptic identity of cINs dictate the differential integration of presynaptic inputs, likely due to expression of specific receptors. In addition, we discovered that specific inputs target PV versus SST cINs at different developmental timepoints. The distinct temporal dynamics when these afferents are established also likely reflects these transcriptional differences. However, while cIN identity may strongly regulate circuit refinement, cINs appear to be minimally altered by their afferents. Specifically, recent transcriptomic data have shown that gene expression in M2 and V1 cINs is very similar ^4^. This is in contrast to the differential gene expression observed in excitatory cells within distinct cortical regions. Recent work suggests that cINs are evolutionary more conserved than neocortical pyramidal cells^73^ supporting a strong intrinsic genetic contribution to their development.

In sensory areas, thalamocortical afferents to cINs appear to follow distinct rules. While inhibition can control thalamocortical pathway development^74^, it has been shown that sensory activity differentially regulates PV circuit inhibition^75,76^ and plasticity^43,77-79^. We demonstrated here that activity impacts the development of PV cINs, while TC pathways onto SST cINs are genetically regulated. In addition, we confirmed our previous observation that SST cINs transiently project to PV cINs and are necessary for the development of feedforward inhibition (FFI) ^12,43^. Interestingly, in M2 the organization of inhibitory circuitry is quite different. In this area, PV cINs maintain transient SST cIN connectivity longer, and receive both FO and HO thalamic afferents at adulthood. Similarly, FFI in mPFC has also been shown to involve HO projections (MD) onto PV cINs^80^. Given these differences, it would be interesting to investigate how FFI develops in associative areas and specifically why SST to PV cIN connections are maintained in these areas in mature animals. An additional element that potentially contributes to the formation of TC-cIN circuits is the subplate. The subplate^81^ is a transient developmental structure that has been shown to control the maturation of GABA receptors^82^ and the development of TC projections within the cortical plate^83^. We observed that the timing of regression of subplate inputs onto PV and SST cINs is both cell type and area specific (Fig4b). This possibly suggests a role for the subplate in differentially regulating TC connectivity to SST and PV cINs within distinct areas.

Given our discovery that TC connectivity onto SST cINs is disrupted in *Fmr1* but not *Shank3b* mutants, further use of rabies to explore aberrations in cIN connectivity in other neural developmental disorders is warranted. The differences in both the developmental and adult afferent connectivity onto PV and SST cINs within sensory versus associative areas are striking. This suggests that circuit abnormalities in neurodevelopmental disorders are both region and cell-type specific. This emphasizes the necessity of understanding circuit components with respect to their organization within specific functional areas.

While currently available drugs broadly target receptors expressed on all neurons, our results suggest the need to target cells imbedded within specific circuits. Although cINs are attractive targets for such manipulations, their genetic similarity across circuits suggests that finding drugs that selectively target those in particular cortical regions will prove difficult. Recently, we identified a regulatory element selective for a subpopulation of PV cINs84. The targeted use of viruses utilizing such elements may provide a promising therapeutic avenue to explore. Regardless of the means, developing tools to target cINs within distinct cortical regions will be essential for correcting both cognition and sensory processing in neuropsychiatric disease.

## Supporting information

ExtendedData

## Acknowledgements

We thank Marian Fernandez-Otero for her support along the project. This work was supported by grants from the National Institutes of Health (NIH): MH071679, NS08297, NS074972, MH095147, as well as support from the Simons Foundation (SFARI) (to G.F); EMBO Long-Term fellowship, Early and advanced Swiss Foundation postdoctoral fellowships, a Hearst foundation grant (to GP).

## Author contributions

G.P and GF conceived the project and wrote the manuscript. GP and CA carried out injection experiments. SK and GP processed brains for quantification. YB designed and performed the automated analysis with the help of GP. GP and AM quantified the connectivity manually. RC contributed to the design of automated analysis. KDR provided guidance an N2cRV and AAV use as well as provided for nuclear-tdtomato N2RV. EF performed the optogenetic recordings.

## Methods

Animals.All experiments were approved by and in accordance with Harvard Medical School IACUC protocol number IS00001269. C57Bl/6 mice were used fo r breeding with transgenic mice. Transgenic mice, PV-cre (stock number: 017320), SST-cre (stock number: 013044), SST-FlpO (stock number: 031629), Lhx6-iCre (stock number: 026555), Fmr1 KO (stock number: 003025), RCE:loxP (expressing eGFP, stock number: 010701), Ai9 (expressing tdTomato, stock number: 007909) are available at Jackson Laboratories. Both female and male were used in the entire study except for the Fmr1 KO and their littermate controls, which were only males.

### Sensory deprivations

To deprive mice from whisker and visual sensory input, infraorbital nerve section (IONS) and enucleation were performed as previously described49. P0 mouse pups were anesthetized by hypothermia. For IONS, a unilateral skin incision was made between the eye and the whisker pad, and the infraorbital nerve, which innervates the whisker pad, was carefully cut with sterile blade. For enucleation, a small incision was made between the eyelids with a scalpel and the eye was separated from the optic nerve with microscissors in order to be removed from the orbit with forceps. The pups were allowed to recover on a heating pad before being returned to their mother.

Histology. Mice at between P42-P46 for the adult time point or P10 for the developmental time point (n= 3 or 4 for each condition) were perfused with 4% paraformaldehyde (PFA) and brains were fixed overnight in 4% PFA at 4?°C. 50-μm vibratome sections were used for all histological experiments. Every 4 section was collected for the representation of each brain and the sections were processed for immunohistochemistry in order to amplify and stabilize the signals and confirm somatostatin identity.

For the immunofluorescence, brain sections were incubated 1 h at room temperature in a blocking solution containing 3% Normal Donkey serum and 0.3% Triton X-100 in PBS and incubated overnight or 2 nights at 4°C with primary antibodies: rat anti-RFP (1:1,000; Chromotek #5f8), chicken anti-GFP (1:1,000; Aves Labs #1020), rabbit anti-somatostatin (1:3,000; Peninsula Laboratories International T-4103.0050), goat anti-ChAT (1:250; Millipore AB144P), goat anti-CTGF (1:500; Santa Cruz Biotechnology sc-14939), rabbit anti-TPH2 (1:500, Novus Biologicals NB74555). Sections were rinsed three times in PBS and incubated for 60–90 min at room temperature or overnight at 4°C with the Alexa Fluor 488-, 594- or 647-conjugated secondary antibodies (1:500; Thermo Fisher Science or Jackson ImmunoResearch).

### Rabies tracing

For adult mice, stereotactic injections were performed between P30-P35. Recombinant AAV-DIO-helpers and N2cRV were diluted at a ratio 1:3 and 23nl were microinjected using NanojectIII at 1nl/second according to stereotaxic coordinates (from Bregma. AP+1.5, ML-0.7, DV+0.85 for M2; AP -1, ML-3, DV+0.89 for S1; AP-3, ML-2.5DV+0.50 for V1). Animals were perfused 9-12 days later. For postnatal time points stereotaxic injections were possible using a neonate adapter (Harvard apparatus). Mouse pups were anesthetized by hypothermia and stereotaxically micro-injected with the rAAV-DIO or fDIO-helpers at P0 and separately with the N2cRV at P5. Animals were perfused 5 days later at P10. All coordinates were determined to target mainly the deeper layer (5-6) of the cortex.

### Viruses

#### rAAV1-DIO-helpers (Cre-ON helpers)

N2cG protein was cloned instead of B19G into the previously published TVA-eGFP construct^85^. This construct was designed with less restriction sites, a small WPRE and human Synapsin promoter to reduce the total length of the genome and for a high neuronal expression.

#### rAAV1-flrtDIO-helpers (Cre-ON, Flp-OFF helpers)

deleting FRT sites were added around the Lox sites of Cre-ON construct, to trigger whole insert deletion upon FlpO expression.

#### rAAV1-hChR2(H134R)-eYFP

Produced from the Addgene plasmid #26973 a gift from K. Deisseroth.

rAAV1-hChR2(H134R)-mCherry. Produced from the Addgene plasmid #26976 a gift from K. Deisseroth.

#### Rabies

EnvA-pseudotyped CVS-N2c(deltaG)- FlpO-mCherry was used. In addition, to simplify automatic detection of cells, we generated Rabies with nuclear expression of reporter: tdTomato with a nuclear localization signal (NLS-tdtomato) was cloned instead of tdTomato into RabV CVS-N2c(deltaG-tdTomato) plasmid previously published^7^ and a gift from T. Jessell as Addgene #73462.

Both rabies types were either produced, amplified and EnvA-pseudotyped in lab, or generously shared by K. Ritola.

### Imaging and Data analysis (yannick’s methods name+ manual counting)

Each brain section containing labelled cells was acquired as a tiled image on a motorized Zeiss Axio Imager A1. Brains with no helper expression were used as control of good pseudotyping and specificity of the EnvA-N2cRV (data not shown). Starter cells (colocalization of GFP^+^ helpers and tdTomato^+^ N2cRV) were manually quantified on Adobe Photoshop software.

Representative brains with less than 10 starter cells were discarded. The highest concentration of starter cells was in deeper layers for all brains (L5-6. Extended Data Fig. 1a).

RFP+ retrogradely labeled cells were registered for each region of the Allen Reference Brain atlas for adult brain and of the “Atlas of Developing Mouse Brain at P6” from George Paxinos 2006.

All statistical analysis were performed using Prism (GraphPad).

### In vitro electrophysiology

Optogenetic was performed after expression of ChR2 (rAAV1-hChR2(H134R)-eYFP or rAAV1-hChR2(H134R)-mCherry in the thalamus of SST-cre:RCE or SST-cre:Ai9 mice using stereotaxic injection (see rabies tracing method) in at P10 for mature time points and at P0 for P7 time point. P19-P23 or P7 mice were deeply anesthetized with isoflurane followed by decapitation. The brain was quickly removed and immersed in ice-cold oxygenated (95% O2 / 5% CO2) sucrose cutting solution containing 119 mM NaCl, 2.5 mM KCl, 1.3 mM MgCl, 2.5 mM CaCl2, 1.0 mM Na2HPO4, 26.2 mM NaHCO3, and 11 mM glucose. 300 μm thick coronal slices were cut using a Leica VT 1000S vibratome through primary somatosensory cortex. Slices recovered in a holding chamber with artificial cerebrospinal fluid (ACSF) containing (in mM) 125 NaCl, 20 Glucose, 2.5 KCl, 1.25 NaH2PO4, 26 NaHCO3, 2 CaCl2, 1 MgCl2 (pH=7.4) at 32 degrees C for 30 minutes and at room temperate for at least 45 minutes prior to recordings. For recordings slices were transferred to an upright microscope (Scientifica) with oblique illumination Olympus optics. Cells were visualized using a 60x water immersion objective. Slices were perfused with ACSF in a recording chamber at 2 ml/min at room temperature. All slice preparation and recording solutions were oxygenated with carbogen gas (95% O2, 5% CO2, pH 7.4). Patch electrodes (3–6 MΩ) were pulled from borosilicate class (1.5 mm OD, Harvard Apparatus). For all recordings patch pipettes were filled with an internal solution containing (in mM): 125 Cs-gluconate, 2 CsCl, 10 HEPES, 1 EGTA, 4 MgATP, 0.3 Na-GTP, 8 Phosphocreatine-Tris, 1 QX-314-Cl, equilibrated with CsOH at pH=7.3. Recordings were performed using a Multiclamp 700B amplifier (Molecular Devices) and digitized using a Digidata 1550A and the Clampex 10 program suite (Molecular Devices). Voltage-clamp signals were filtered at 3 kHz and recorded with a sampling rate of 10 kHz. Recordings were performed at a holding potential of -70 mV. Cells were only accepted for analysis if the initial series resistance was less than 40 MΩ and did not change by more than 20% during the recording period. The series resistance was compensated at least ∼50% in voltage-clamp mode and no correction was made for the liquid junction potential.

Under low magnification, the barrels could be readily identified allowing to target the deeper Layer 5 of the cortex and high-power magnification was used to guide the recording electrode onto visually identified neurons. Whole-cell patch-clamp recordings were obtained from tdTomato-expressing SST cINs or adjacent non-fluorescent pyramidal-shaped neurons in L5. The following number of cells were recorded for each condition: P7: 10 SST, 4 pyramidal; P19-21: 14 SST, 6 pyramidal.

To activate thalamic afferents expressing ChR2, blue light was transmitted from a collimated LED (Mightex) attached to the epifluorescence port of the upright microscope. 1 ms pulses of light were directed to the slice in the recording chamber via a mirror coupled to the 60x objective (1.0 numerical aperture). Flashes were delivered every 5 s for a total of 15 trials. The LED output was driven by a transistor-transistor logic output from the Clampex software.

Recordings were performed in the presence of 1 µm TTX and 1 mM 4-AP (Tocris).

Data analysis was performed off-line using the Clampfit module of pClamp (Molecular Devices) and Prism 8 (GraphPad). The amplitude of evoked synaptic currents was obtained by averaging the peak amplitude of individual waveforms over 15 trials per cell.

EPSC amplitudes recorded from SST interneurons were then normalized for injection size by dividing the average EPSC by the evoked current amplitude from a putative L5b pyramidal neuron in the same animal.

#### Legends

**Extended Data Fig. 1: layer distribution, normalization and Cre lines. a**, Deep layer cortical distribution of the starter cells for each n grouped per time point and area. squares = PV starter cells, circles = SST starter cells. **b**, relative numbers of total retrogradely labeled neurons targeting either PV or SST cINs within M2, S1 and V1, in adult and development. **c**, Onset of reporter expression upon SST-Cre and PV-cre drivers. scale bar 100μm. **d**, Percentage of somatostatin negative (PV cINs) vs somatostatin positive in AAV-IS-helpers infected population at P5. PV cINs 73.67% ±3.03 in M2; 73.77%±0.84 in S1; 76.60±3.96 in V1.

**Extended Data Fig. 2: Area-specific connectivity a**-**d**, Degree of connectivity to PV and SST cINs in M2, S1 and V1 from ipsilateral cortical projections (one-way ANOVA, p<0.01) (**a**), from the claustrum region of the cortex (one-way ANOVA, p<0.01) (**b**), from amygdala (one-way ANOVA, p<0.05) (**c**), from hypothalamus (**d**). **e**, Cholinergic identity of rabies labeled basal forebrain neurons (red), with ChAT staining (green). Scale bar 100μm. **f**, Network of all presynaptic structures projecting to PV and SST cINs in M2, S1 and V1 using the Fruchterman-Reingold algorithm arrange cINs along their areal identity.

**Extended Data Fig. 3: Cortical projections to PV and SST cINs are changed over development (1) a**-**d**, Degree of connectivity to PV and SST cINs in M2, S1 and V1 both in adult and developmental timepoints from local cortex (**a**), contralateral cortex (**b**), ipsilateral cortex (**c**), subplate (**d**). **e**, SST cINs projections to PV cINs in M2, S1 and V1 in adult and during development. **f**, (left) Example of somatostatin staining (blue) onto retrograde N2cRV cells (red); (right) Subplate identity of N2cRV deep cortical neurons (red) using CTGF staining (blue). scale bar 50μm.

**Extended Data Fig. 4: Afferents to PV and SST cINs are changed over development (1)** Degree of connectivity to PV and SST cINs in M2, S1 and V1 both in adult and developmental timepoints from amygdala (**a**), hypothalamus (**b**), medial septum of the basal forebrain (**c**), diagonal band of the basal forebrain (**d**), ventral pallidum and substantia innominate of the basal forebrain (**e**), Mindbrain and Hindbrain (**f**), dopaminergic structures (VTA, STN, ZI, FF) (**g**), claustrum (**h**).

**Extended Data Fig. 5: Thalamocortical connectivity to SST and PV cINs a**, FO, HO and limbic identification of all thalamocortical nuclei (at rostral and caudal anatomical levels) as previously described^48,49^. **b**, Heatmap representing the percentage of thalamic connectivity onto PV and SST in M2, S1 and V1, as well as example of tracing. Scale bar 200μm. **c**, Degree of connectivity to PV and SST cINs in M2, S1 and V1 from the whole thalamus (**c**), from the FO and HO neurons of the thalamus (**d**).

**Extended Data Fig. 6: Thalamocortical projections to S1 SST cINs in whisker-deprived animals a-d**, Degree of connectivity from the whole thalamus (**a**) and FO/HO/limbic neurons (**b**) to PV cINs in Ctrl and sensory deprived animals (IONS) and to SST cINs (**c** and **d**).

**Extended Data Fig. 7: Thalamocortical projections to S1 SST cINs in Fmr1 knock-out mice a-b**, Degree of connectivity from the whole thalamus (**a**) and FO/HO/limbic neurons (**b**) to SST cINs.

## References

1. Barthas, F. & Kwan, A. C. Secondary Motor Cortex: Where ‘Sensory’ Meets ‘Motor’ in the Rodent Frontal Cortex. Trends Neurosci (2016). doi:10.1016/j.tins.2016.11.006

2. Manita, S. et al. A Top-Down Cortical Circuit for Accurate Sensory Perception. NEURON 86, 1304–1316 (2015).

3. Muñoz, W. & Rudy, B. Spatiotemporal specificity in cholinergic control of neocortical function. Current Opinion in Neurobiology 26, 149–160 (2014).

4. Pouchelon, G. & Jabaudon, D. Nurturing the cortex’s thalamic nature. Current Opinion in Neurology 1 (2014). doi:10.1097/WCO.0000000000000070

5. Chou, S.-J. et al. Geniculocortical input drives genetic distinctions between primary and higher-order visual areas. Science 340, 1239–1242 (2013).

6. Li, H. et al. Laminar and columnar development of barrel cortex relies on thalamocortical neurotransmission. NEURON 79, 970–986 (2013).

7. Tasic, B. et al. Shared and distinct transcriptomic cell types across neocortical areas. Nature 563, 72–78 (2018).

8. Reardon, T. R. et al. Rabies Virus CVS-N2c(ΔG) Strain Enhances Retrograde Synaptic Transfer and Neuronal Viability. NEURON (2016). doi:10.1016/j.neuron.2016.01.004

9. Wall, N. R. et al. Brain-Wide Maps of Synaptic Input to Cortical Interneurons. Journal of Neuroscience 36, 4000–4009 (2016).

10. Zhang, S. et al. Organization of longrange inputs and outputs of frontal cortex for top-down control. Nat Neurosci 19, 1733–1742 (2016).

11. Ährlund-Richter, S. et al. A wholebrain atlas of monosynaptic input targeting four different cell types in the medial prefrontal cortex of the mouse. Nat Neurosci 22, 657–668 (2019).

12. Sun, Q. et al. A whole-brain map of long-range inputs to GABAergic interneurons in the mouse medial prefrontal cortex. Nat Neurosci 1 (2019). doi:10.1038/s41593-019-0429-9

13. Clascá, F., Rubio-Garrido, P. & Jabaudon, D. Unveiling the diversity of thalamocortical neuron subtypes. Eur J Neurosci 35, 1524–1532 (2012).

14. Sauerbrei, B. A. et al. Cortical pattern generation during dexterous movement is input-driven. Nature 1–6 (2019). doi:10.1038/s41586-019-1869-9

15. Anastasiades, P. G., Marlin, J. J. & Carter, A. G. Cell-Type Specificity of Callosally Evoked Excitation and Feedforward Inhibition in the Prefrontal Cortex. Cell Rep 22, 679–692 (2018).

16. Fmr1 knockout mice: a model to study fragile X mental retardation. The Dutch-Belgian Fragile X Consortium. Cell 78, 23–33 (1994).

17. Gonçalves, J. T., Anstey, J. E., Golshani, P. & Portera-Cailliau, C. Circuit level defects in the developing neocortex of Fragile X mice. Nat Neurosci 16, 903–909 (2013).

18. Kohara, K. et al. Cell type-specific genetic and optogenetic tools reveal hippocampal CA2 circuits. Nat Neurosci 17, 269–279 (2014).

19. Fenno, L. E. et al. Targeting cells with single vectors using multiple-feature Boolean logic. Nat Meth 11, 763–772 (2014).

20. Zhang, S. et al. Whole-Brain Mapping of Monosynaptic Afferent Inputs to Cortical CRH Neurons. Front. Neurosci. 13, 565 (2019).

21. Porter, J. T., Johnson, C. K. & Agmon, A. Diverse types of interneurons generate thalamus-evoked feedforward inhibition in the mouse barrel cortex. Journal of Neuroscience 21, 2699–2710 (2001).

22. Silberberg, G. & Markram, H. Disynaptic inhibition between neocortical pyramidal cells mediated by Martinotti cells. NEURON 53, 735–746 (2007).

23. Cummings, K. A. & Clem, R. L. Prefrontal somatostatin interneurons encode fear memory. Nat Neurosci 23, 61–74 (2020).

24. Distinct Inhibitory Circuits Orchestrate Cortical beta and gamma Band Oscillations. NEURON 96, 1403–1418.e6 (2017).

25. Yu, J., Hu, H., Agmon, A. & Svoboda, K. Recruitment of GABAergic Interneurons in the Barrel Cortex during Active Tactile Behavior. NEURON (2019). doi:10.1016/j.neuron.2019.07.027

26. Miller, F. D. & Gauthier, A. S. Timing is everything: making neurons versus glia in the developing cortex. NEURON 54, 357–369 (2007).

27. Paton, J. J. & Buonomano, D. V. The Neural Basis of Timing: Distributed Mechanisms for Diverse Functions. NEURON 98, 687–705 (2018).

28. Kanold, P. O. & Luhmann, H. J. The subplate and early cortical circuits. Annu. Rev. Neurosci. 33, 23–48 (2010).

29. Kanold, P. O., Kara, P., Reid, R. C. & Shatz, C. J. Role of subplate neurons in functional maturation of visual cortical columns. Science 301, 521–525 (2003).

30. Hoerder-Suabedissen, A. et al. Novel markers reveal subpopulations of subplate neurons in the murine cerebral cortex. Cerebral Cortex 19, 1738–1750 (2009).

31. Tuncdemir, S. N. et al. Early Somatostatin Interneuron Connectivity Mediates the Maturation of Deep Layer Cortical Circuits. NEURON 89, 521–535 (2016).

32. Pouchelon, G. et al. Modality-specific thalamocortical inputs instruct the identity of postsynaptic L4 neurons. Nature 511, 471–474 (2014).

33. Katz, L. C. & Shatz, C. J. Synaptic activity and the construction of cortical circuits. Science 274, 1133–1138 (1996).

34. Fox, K. & Wong, R. O. L. A comparison of experience-dependent plasticity in the visual and somatosensory systems. NEURON 48, 465–477 (2005).

35. Hubel, D. H. & Wiesel, T. N. The period of susceptibility to the physiological effects of unilateral eye closure in kittens. The Journal of Physiology 206, 419–436 (1970).

36. Maffei, A., Nataraj, K., Nelson, S. B. & Turrigiano, G. G. Potentiation of cortical inhibition by visual deprivation. Nature 443, 81–84 (2006).

37. Miska, N. J., Richter, L. M., Cary, B. A., Gjorgjieva, J. & Turrigiano, G. G. Sensory experience inversely regulates feedforward and feedback excitation-inhibition ratio in rodent visual cortex. Elife 7, (2018).

38. Antón-Bolaños, N., Espinosa, A. & López-Bendito, G. Developmental interactions between thalamus and cortex: a true love reciprocal story. Current Opinion in Neurobiology 52, 33–41 (2018).

39. Chittajallu, R. & Isaac, J. T. R. Emergence of cortical inhibition by coordinated sensory-driven plasticity at distinct synaptic loci. Nat Neurosci 13, 1240–1248 (2010).

40. Chattopadhyaya, B. et al. Experience and activity-dependent maturation of perisomatic GABAergic innervation in primary visual cortex during a postnatal critical period. Journal of Neuroscience 24, 9598–9611 (2004).

41. Hensch, T. K. & Stryker, M. P. Columnar architecture sculpted by GABA circuits in developing cat visual cortex. Science 303, 1678–1681 (2004).

42. Phillips, J. W. et al. A repeated molecular architecture across thalamic pathways. Nat Neurosci 22, 1925–1935 (2019).

43. Frangeul, L. et al. A cross-modal genetic framework for the development and plasticity of sensory pathways. Nature (2016). doi:10.1038/nature19770

44. Ackman, J. B., Burbridge, T. J. & Crair, M. C. Retinal waves coordinate patterned activity throughout the developing visual system. Nature 490, 219–225 (2012).

45. Burbridge, T. J. et al. Visual circuit development requires patterned activity mediated by retinal acetylcholine receptors. NEURON 84, 1049–1064 (2014).

46. Marín, O. Interneuron dysfunction in psychiatric disorders. Nat Rev Neurosci 13, 107–120 (2012).

47. Schuetze, M. et al. Morphological Alterations in the Thalamus, Striatum, and Pallidum in Autism Spectrum Disorder. Neuropsychopharmacology 41, 2627–2637 (2016).

48. Peñagarikano, O., Mulle, J. G. & Warren, S. T. The pathophysiology of fragile x syndrome. Annu Rev Genomics Hum Genet 8, 109–129 (2007).

49. Oh, S. W. et al. A mesoscale connectome of the mouse brain. Nature 508, 207–214 (2014).

50. Cruikshank, S. J., Urabe, H., Nurmikko, A. V. & Connors, B. W. Pathway-specific feedforward circuits between thalamus and neocortex revealed by selective optical stimulation of axons. NEURON 65, 230–245 (2010).

51. Audette, N. J., Urban-Ciecko, J., Matsushita, M. & Barth, A. L. POm Thalamocortical Input Drives Layer-Specific Microcircuits in Somatosensory Cortex. Cerebral Cortex 1–17 (2017). doi:10.1093/cercor/bhx044

52. Sermet, B. S. et al. Pathway-, layer- and cell-type-specific thalamic input to mouse barrel cortex. Elife 8, (2019).

53. Zhu, X. et al. Rabies Virus Pseudotyped with CVS-N2C Glycoprotein as a Powerful Tool for Retrograde Neuronal Network Tracing. Neurosci Bull 36, 202–216 (2020).

54. Wall, N. R., La Parra, De, M., Callaway, E. M. & Kreitzer, A. C. Differential innervation of direct- and indirect-pathway striatal projection neurons. NEURON 79, 347–360 (2013).

55. Kim, T. et al. Cortically projecting basal forebrain parvalbumin neurons regulate cortical gamma band oscillations. Proceedings of the National Academy of Sciences 112, 3535–3540 (2015).

56. Espinosa, N. et al. Basal Forebrain Gating by Somatostatin Neurons Drives Prefrontal Cortical Activity. Cerebral Cortex 1–12 (2017). doi:10.1093/cercor/bhx302

57. Celada, P., Puig, M. V. & Artigas, F. Serotonin modulation of cortical neurons and networks. Front Integr Neurosci 7, 25 (2013).

58. Szőnyi, A. et al. The ascending median raphe projections are mainly glutamatergic in the mouse forebrain. Brain Struct Funct 221, 735–751 (2016).

59. Gu, Z. & Yakel, J. L. Timing-dependent septal cholinergic induction of dynamic hippocampal synaptic plasticity. NEURON 71, 155–165 (2011).

60. Yaeger, C. E., Ringach, D. L. & Trachtenberg, J. T. Neuromodulatory control of localized dendritic spiking in critical period cortex. Nature 1 (2019). doi:10.1038/s41586-019-0963-3

61. Ballinger, E. C., Ananth, M., Talmage, D. A. & Role, L. W. Basal Forebrain Cholinergic Circuits and Signaling in Cognition and Cognitive Decline. NEURON 91, 1199–1218 (2016).

62. Marques-Smith, A. et al. A Transient Translaminar GABAergic Interneuron Circuit Connects Thalamocortical Recipient Layers in Neonatal Somatosensory Cortex. NEURON 89, 536–549 (2016).

63. Guillery, R. W. & Sherman, S. M. Branched thalamic afferents: what are the messages that they relay to the cortex? Brain Res Rev 66, 205–219 (2011).

64. Uesaka, N., Hayano, Y., Yamada, A. & Yamamoto, N. Interplay between laminar specificity and activity-dependent mechanisms of thalamocortical axon branching. J Neurosci 27, 5215–5223 (2007).

65. Delevich, K., Tucciarone, J., Huang, Z. J. & Li, B. The Mediodorsal Thalamus Drives Feedforward Inhibition in the Anterior Cingulate Cortex via Parvalbumin Interneurons. Journal of Neuroscience 35, 5743–5753 (2015).

66. Vormstein-Schneider, D. et al. Viral manipulation of functionally distinct interneurons in mice, non-human primates and humans. Nat Neurosci 50, 825–8 (2020).

