## ExtendedData for "The organization and developmental establishment of cortical interneuron presynaptic circuits"

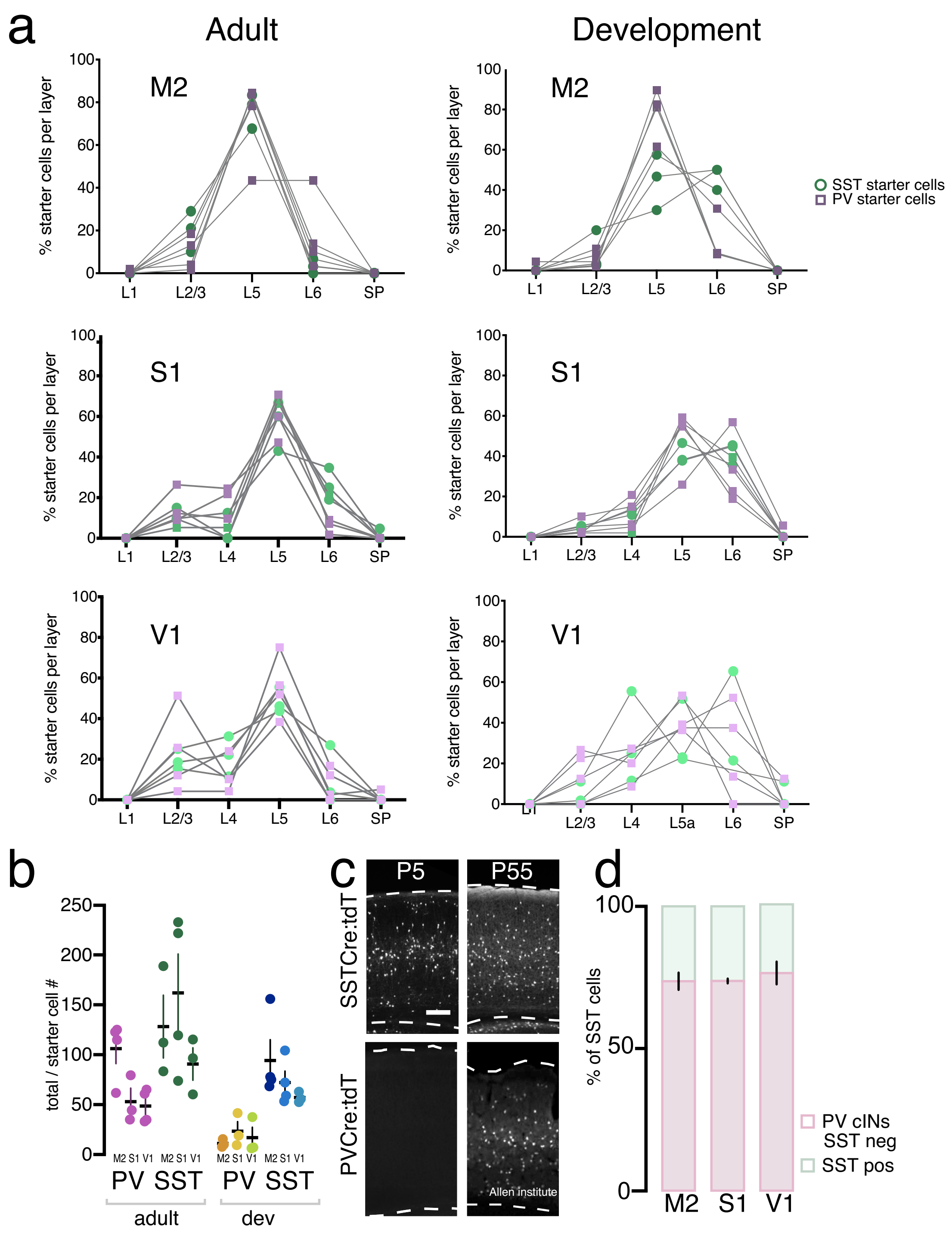


**Extended Data Fig. 1: layer distribution, normalization and Cre lines.**

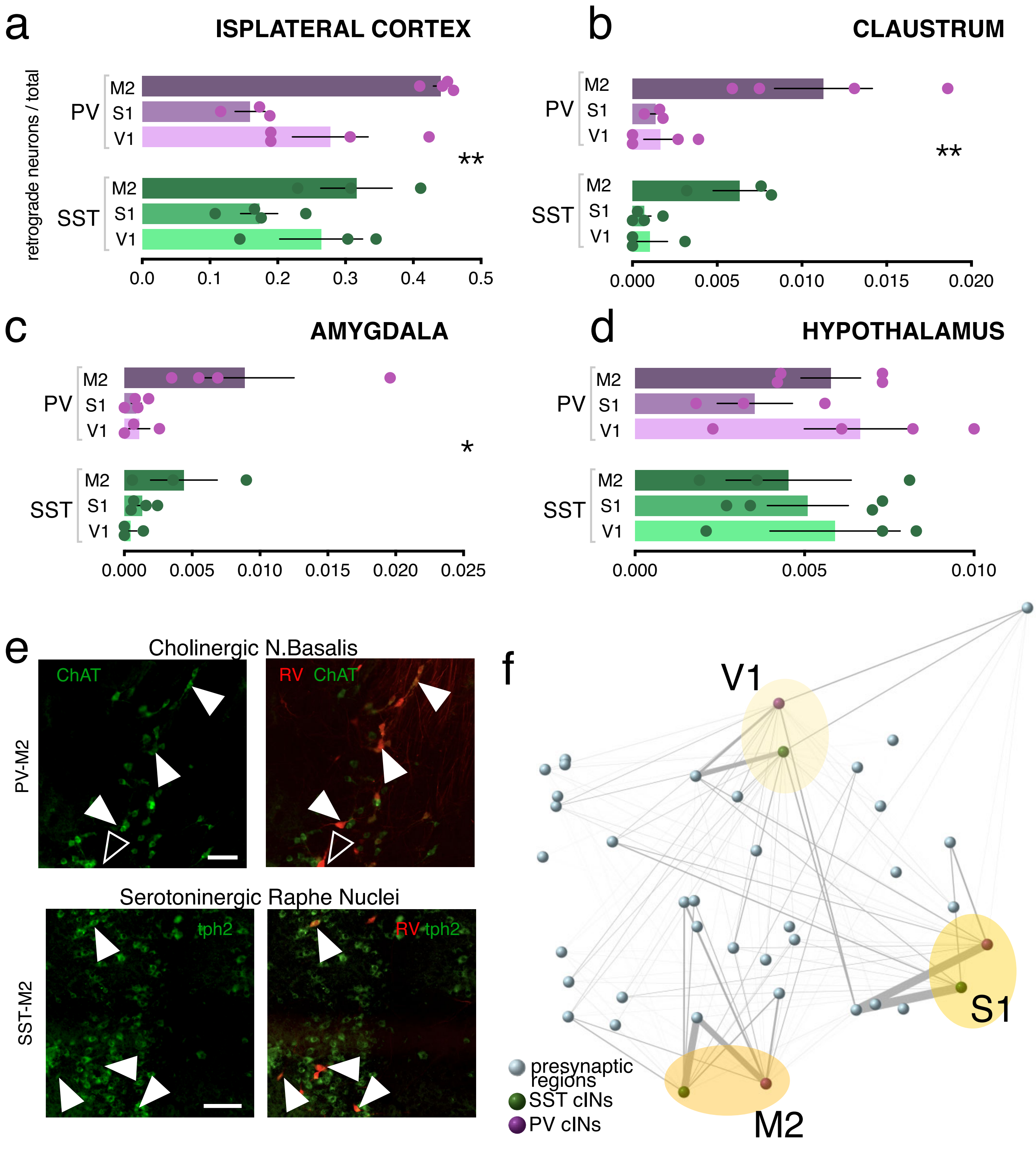


**Extended Data Fig. 2:Area-specific connectivity**

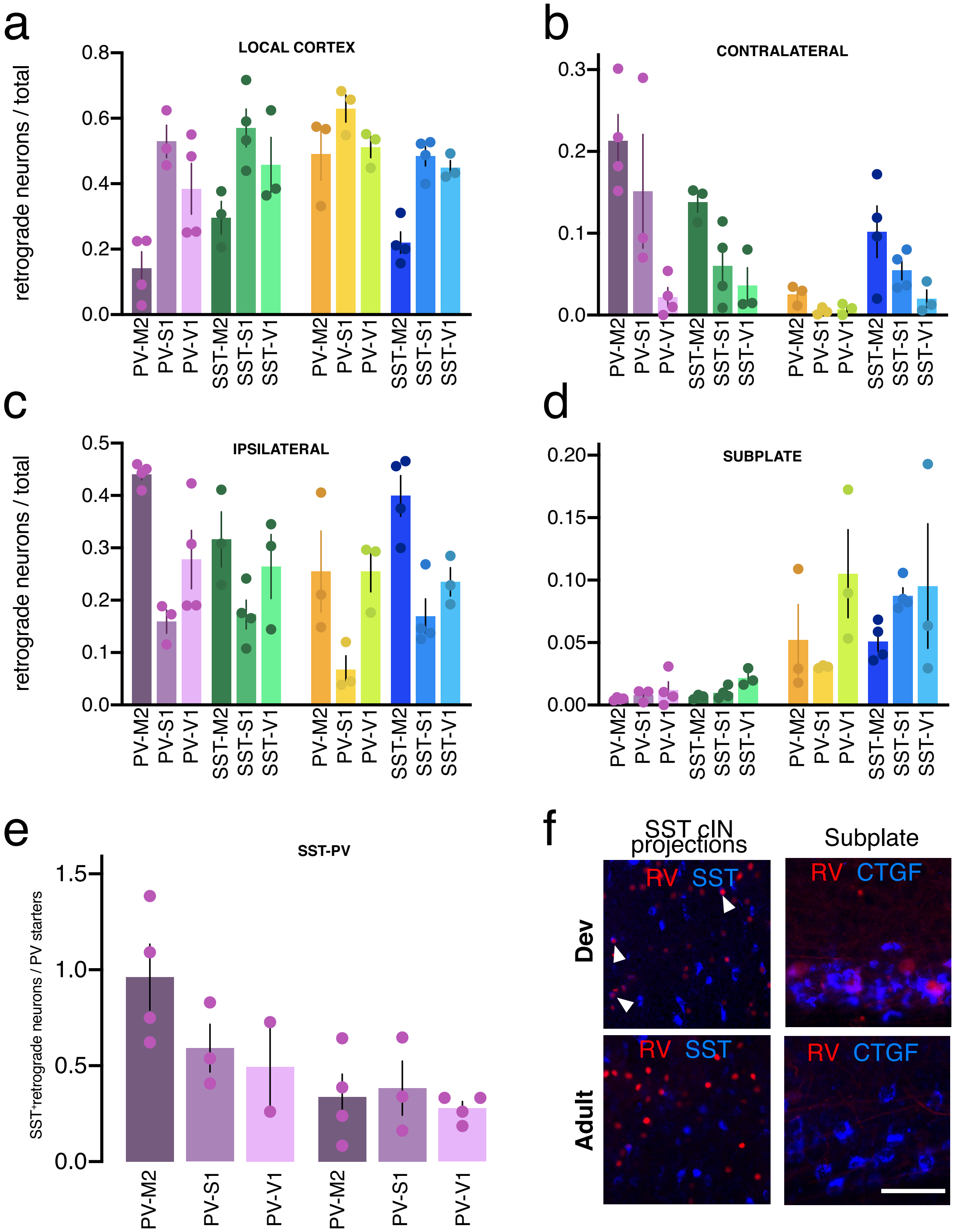


**Extended Data Fig. 3: Cortical projections to PV and SST cINs are changed over development (1)**

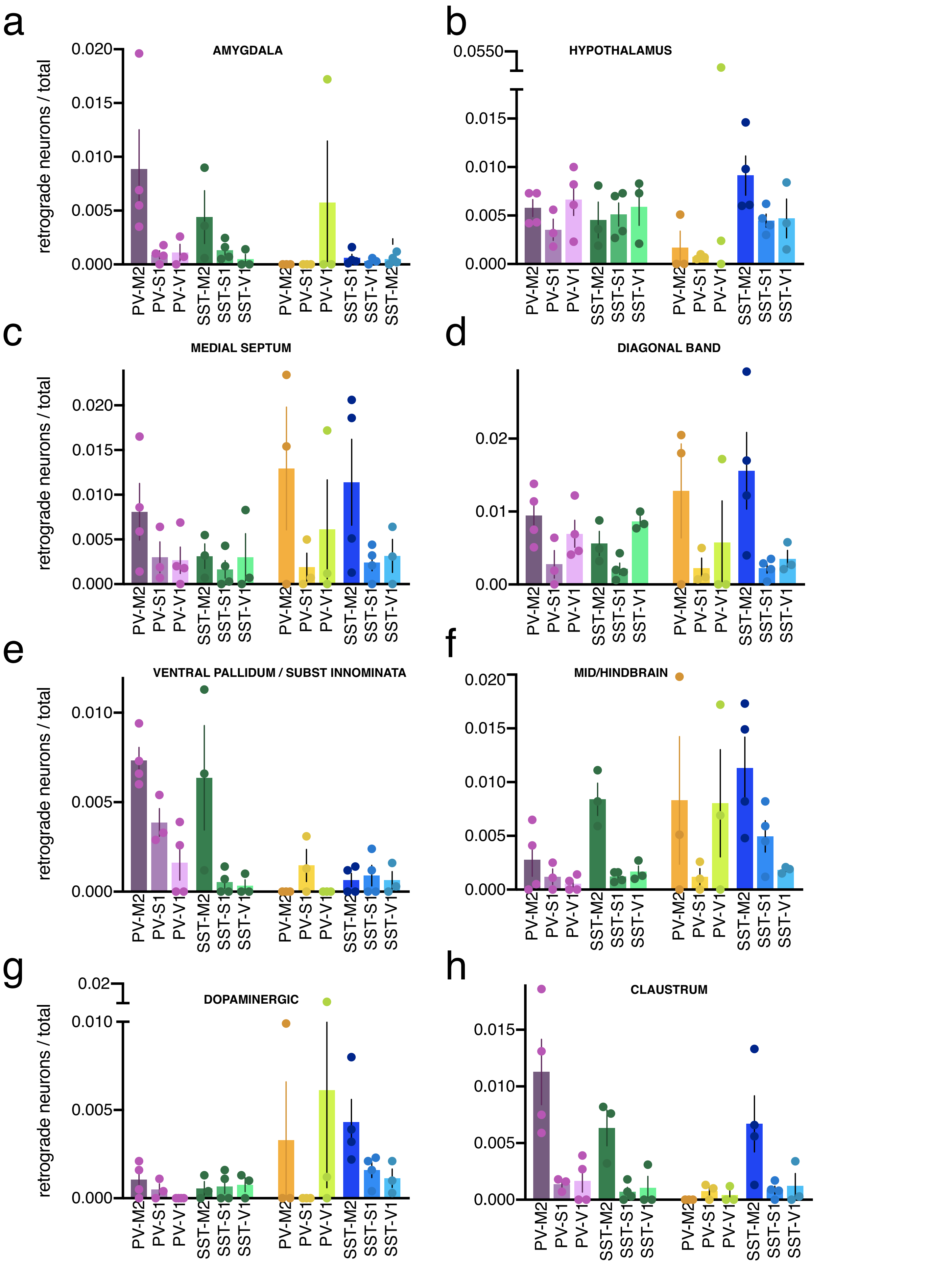


**Extended Data Fig. 4: Afferents to PV and SST cINs are changed over development (2)**

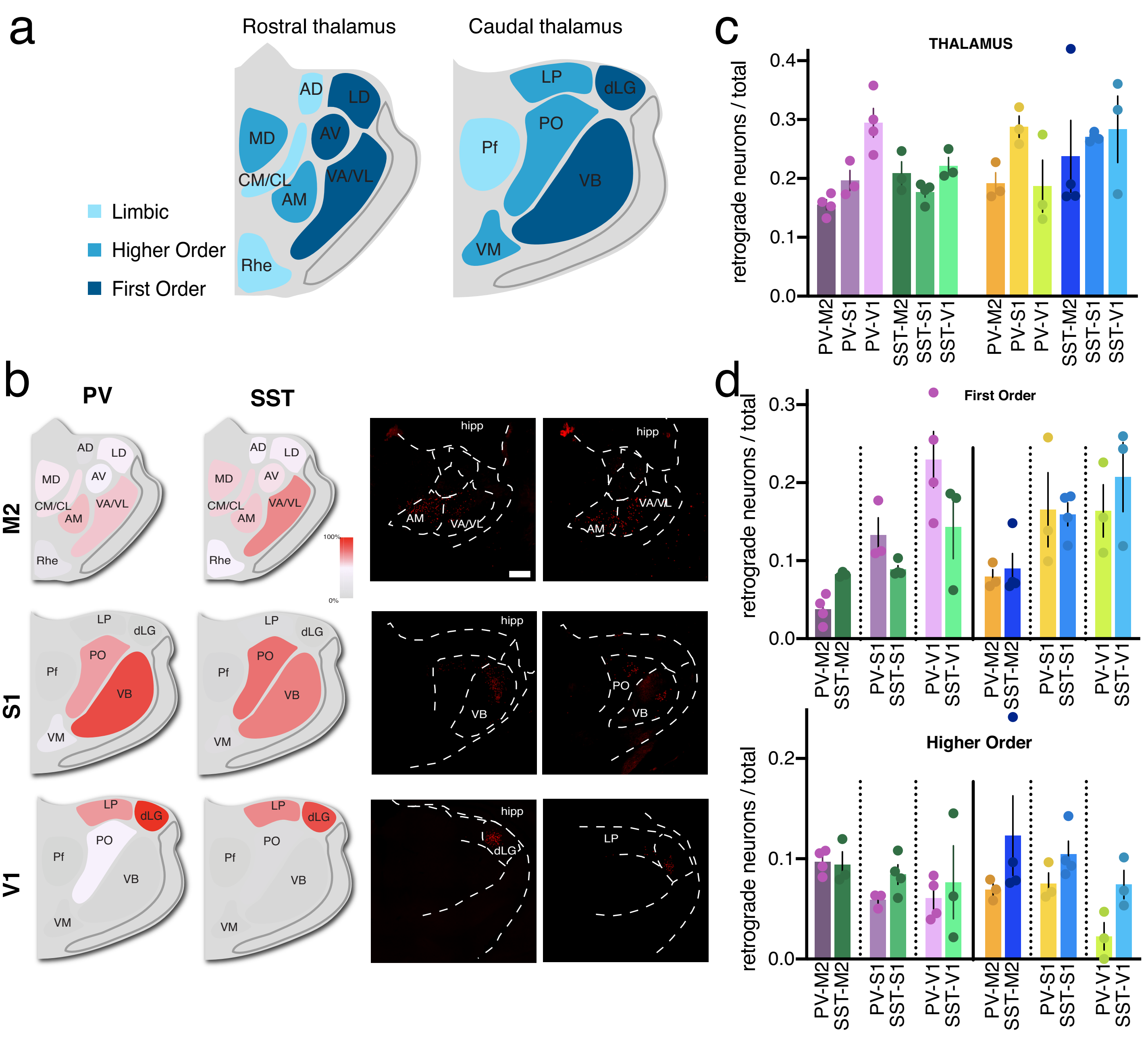


### Extended Data Fig. 5: Thalamocortical connectivity to SST and PV cINs

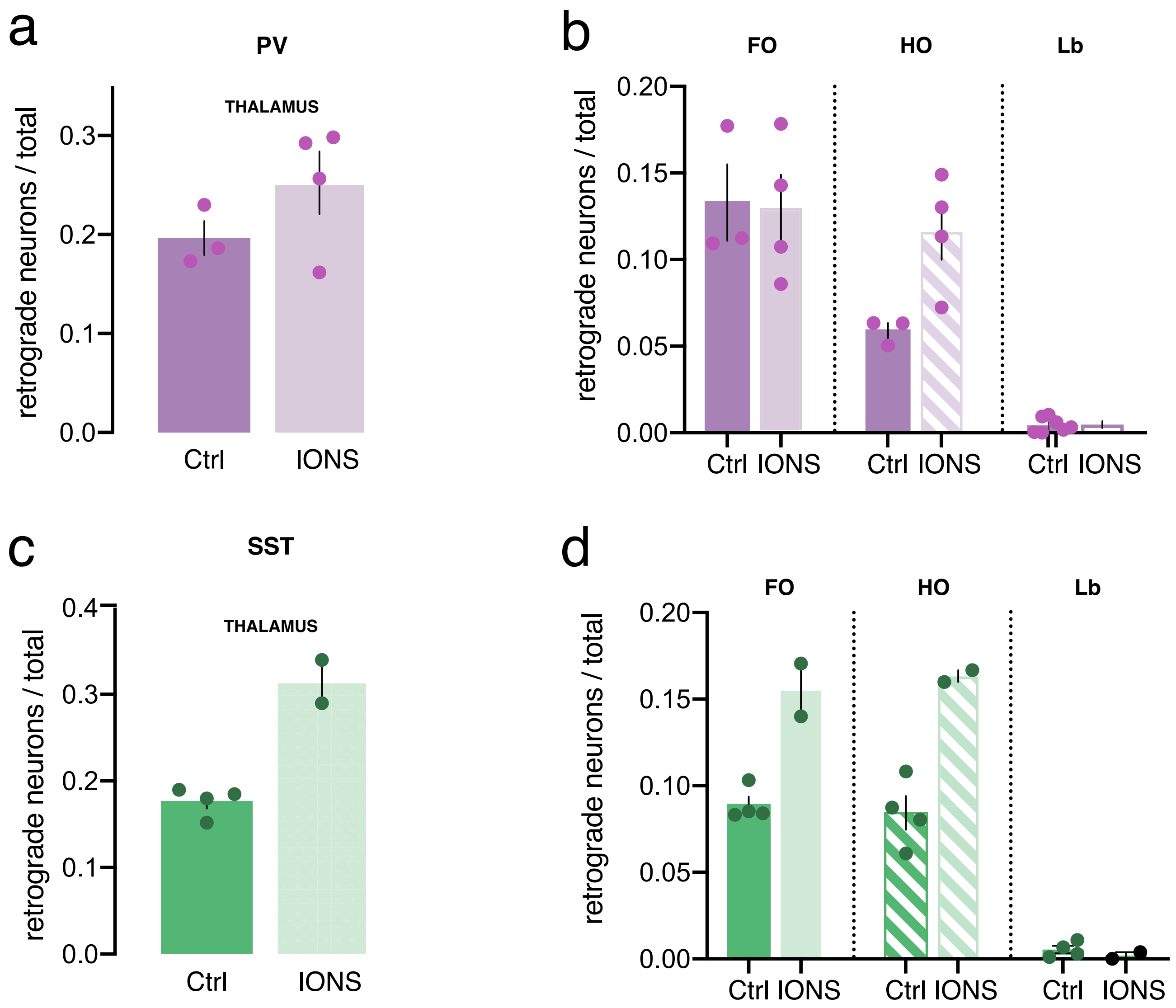


**Extended Data Fig. 6: Thalamocortical projections to S1 SST cINs in whisker-deprived animals**

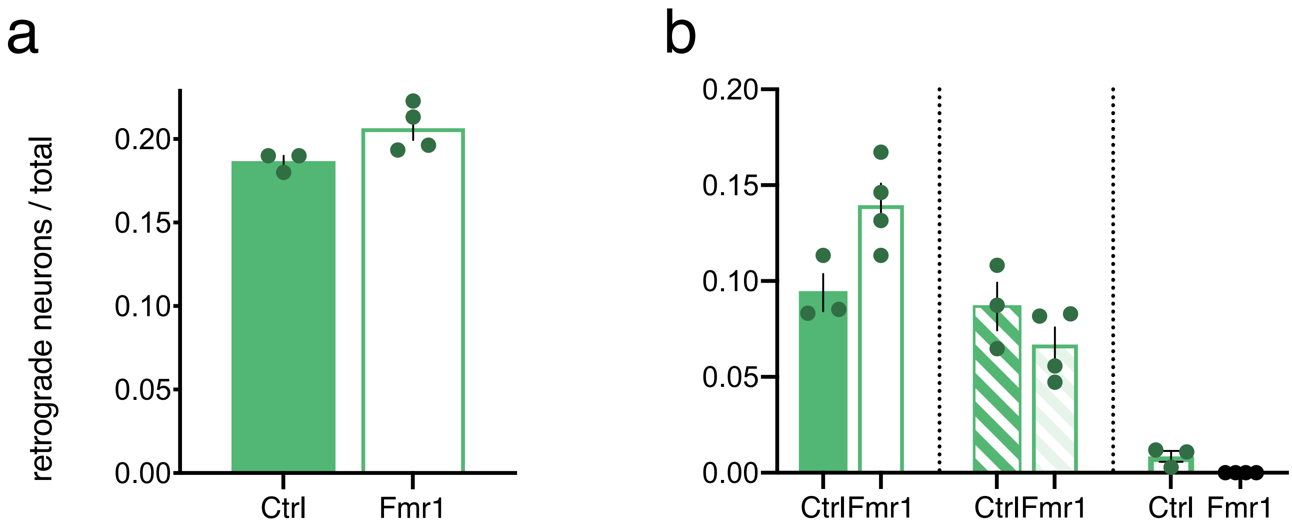


**Extended Data Fig. 7: Thalamocortical projections to S1 SST cINs of Fmr1 knock out mice**
